# Oral administration of engineered recombinant outer membrane vesicles-based nano-vaccine formulation triggers robust mucosal and systemic humoral immunity against poultry pathogens

**DOI:** 10.1101/2025.10.13.682241

**Authors:** Srinivas Duvvada, Farhan Ahmed, Rafiq Ahmad Khan, Sai Nikhith Cholleti, Saima Naz, Mohd Shiraz, Akif Mohd, Tanmay Jana, Madhuri Subbiah, Nooruddin Khan

## Abstract

Although vaccination is an integral part of poultry health management, affordable, scalable, and easily administrable new-generation vaccine discovery platforms are required to combat emerging infectious diseases. Oral vaccines offer an alternative to injectable options owing to their ease of administration; however, existing platforms fail to confer robust protective systemic and mucosal immunity. Here, we used engineered outer membrane vesicles (rOMVs) displaying the desired vaccine antigens, and derived from hypervesiculating probiotic E coli Nissle 1917 (EcN) ^[36]^, which elicits both mucosal and systemic immunity when administered orally. Using such approach, we attempted to formulate rOMV based single Oral vaccine against Newcastle Disease Virus (NDV) and Infectious Bursal Disease Virus (IBDV), which pose significant risks to the poultry sector. We engineered rOMVs to display the antigenic regions of immunodominant Hemagglutinin-Neuraminidase (HN) and Viral Protein 2 (VP2) of NDV and IBDV, respectively, on their surface, with sizes less than 150 nm and considerable polydispersity. The Oral administration of these rOMVs expressing the HN and VP2 induced a robust immune response which shows enhanced production of high-titer antigen-specific sIgA and IgG, which were able to neutralize the virus effectively in an in-vitro infection model system. While the poultry pathogens such as NDV and IBDV served as a model, the rOMV-based platforms hold considerable potential for the development of a multivalent oral biotherapeutic agent, including vaccines against a wide range of emerging human and animal pathogens by engineering rOMVs to display immunodominant antigenic portions, demonstrating its capability as a next-generation oral vaccine discovery platform.

## Introduction

The discovery of vaccines marks a crucial milestone in human history, greatly lowering mortality rates caused by infectious diseases Although wide variety of vaccines discovery platforms are available efforts are being made to develop new generation platforms which is affordable, scalable, efficacious, and safe. Oral vaccine delivery platform offers several advantages over traditional injection-based methods as it is non-invasive and can enhance accessibility, especially in developing and underdeveloped regions where needle-based vaccine administration can be difficult, time-consuming, and necessitate skilled personnel. Using outer Membrane Vesicles (OMVs) as carriers for oral vaccines presents a compelling alternative with numerous benefits ^[1]^. Probiotic OMVs are cost-effective and easy to mass-produce, ensuring their availability and scalability. Furthermore, they hold promise for both preventive and therapeutic uses due to their capacity to elicit long-lasting immune responses ^[2],[3]^. Various studies have highlighted the significant potential of OMVs derived from Shigella, Vibrio cholerae, Salmonella, and other pathogens as potent vaccine candidates, with promising results from preclinical trials ^[4]^ .

OMVs are nanostructure and pathogen mimetic produced by Gram-negative bacteria. They are emerging as promising oral vaccine delivery tool as they can carry the desired biotherapeutic molecules. Naturally OMVs carry wide varieties of biomolecules, including nucleic acids, lipopolysaccharides, and proteins, which are encapsulated in vesicles during bacterial growth and stress ^[5]^. These varied cargoes may trigger a strong immune response. Owing to their unique composition, OMVs can mimic the structure of pathogens and elicit immune responses comparable to those triggered by genuine infections ^[6], [7]^. These properties make OMVs ideal for mucosal immunization because they efficiently deliver antigens to mucosal surfaces, resulting in local and systemic immunity ^[8]^ ^[9]^ ^[10]^. Additionally, OMVs can be engineered to deliver specific antigens, thereby enhancing vaccine efficacy and precision ^[11]^ ^[12]^ ^[13]^ ^[14]^.The development of an oral vaccine for poultry health management is an urgent necessity, as it presents a highly effective and practical alternative to conventional injectable methods. In the agricultural sector, poultry is integral in supplying a significant portion of meat and eggs, thereby contributing substantially to the animal protein intake essential for human health. Globally, the poultry industry is a billion-dollar market, and Asia has one of the largest value shares for meeting global needs^[15]^. As the population is expanding continuously, the demand for poultry products is also increasing, which is outpacing the supply. Poultry meat, especially chicken, is the most commonly consumed lean protein owing to its nutritional value and affordability. Eggs are another key resource that offers rich nutrients and complements various dietary options. It not only provides daily bread for millions of people, but also a source of employment for many people. However, the poultry industry faces remarkable hurdles in its critical role^[16]^. Traditional approaches towards disease control, such as strict biosecurity, use of antibiotics, elimination of infected birds, and vaccination, have been vital in containing outbreaks^[16]^ ^[17]^ ^[18]^. Infectious diseases pose a major threat and can inflict catastrophic damage. Infectious Newcastle disease virus (NDV), Bursal Disease Virus (IBDV), and Infectious Bronchitis Virus (IBV) rapidly spread through flocks and cause widespread death, reduced production, and severe economic consequences ^[19]^. These outbreaks result in annual losses of billions of dollars.

Newcastle disease (ND) is a significant viral infection caused by virulent strains of *Avian Paramyxovirus 1* (APMV1), a single-stranded, non-segmented, negative-sense RNA virus. This virus is classified under the *Paramyxoviridae* family and includes ten recognized serotypes, labeled APMV-1 through APMV-10^[20]^. Infectious bursal disease (IBD) is a highly contagious illness that primarily affects young chickens, leading to immunosuppression^[20]^ ^[21]^. It is caused by the *infectious bursal disease virus* (IBDV), a pathogen of considerable economic concern in the global poultry industry. The development of more aggressive pathogen strains and the growing resistance to current vaccines highlights the urgent requirement for cutting-edge vaccination techniques^[5]^ ^[22]^.

This current investigation explored the immunogenic potential of engineered probiotic outer membrane vesicles (OMVs) as a non-parenteral vaccine delivery platform for inducing robust mucosal immunity. Specifically, we functionalized OMVs derived from probiotic bacteria to present key viral antigens: the hemagglutinin-neuraminidase (HN) protein of Newcastle Disease Virus (NDV) and the VP2 capsid protein of Infectious Bursal Disease Virus (IBDV). This approach leverages the inherent biocompatibility and stability of OMVs within the gastrointestinal environment to facilitate targeted antigen presentation to the mucosal immune inductive sites^[23]^ ^[24]^. The nanoscale dimensions and intrinsic immunomodulatory properties of OMVs promote efficient uptake by antigen-presenting cells within the gut-associated lymphoid tissue (GALT), thereby initiating both local and systemic immune responses^[25]^. The production of secretory IgA antibodies at mucosal sites and the development of cellular immunity offers a significant advantage in preventing infections at their site of entry. The potential of rOMVs has been demonstrated using poultry pathogens including NDV and IBDV; however, it holds considerable promise for the development of multivalent oral biotherapeutic agents including vaccines targeting a wide array of emerging pathogens, thereby underscoring the efficacy of rOMVs as a next-generation platform for oral vaccine discovery for the treatment of human and animal infectious diseases.

## Materials and Methods

### Synthesis and cloning of genes encoding VP2 and HN in the pET23a vector

Antigenic sequences of HN and VP2 genes ^[26]^ ^[27]^ were chemically synthesized and were procured from Biotech Desk Pvt. Ltd. These genes were cloned into the pET23a vector between BamH1 and Sac1 restriction sites. The clones were further confirmed through BamH1 and SacI-mediated double digestion. Both the clones generated a fragment corresponding to the actual insert upon double digestion. Insert sizes of 378bp (VP2) and 432bp (HN) were obtained upon double digestion. Restriction digestion using BamH1 and Sac1 of Clones for confirmation of VP2 and HN genes in the pET23a vector. Insert size of the genes is indicated in Figure S1.

### Expression and purification of recombinant HN and VP2 proteins of NDV and IBDV respectively

We started with the expression and purification of the HN protein of NDV and VP2 protein of IBDV. The cloned construct HN-pET23a and VP2-pET23a were transformed into *E.coli* Rosetta cells to check the expression of the clone. Briefly, a single colony from the transformed plate was inoculated for the primary culture in Chloramphenicol (35µg/ml), and kanamycin (50µg/ml) containing Luria Bertani (LB) broth. Inoculated culture was incubated overnight in a shaking incubator. One percent of the primary culture was used as the starting material for the secondary culture. The secondary culture was grown in an incubator with shaking, at 37°C until an optical density (OD) of 0.5-0.7 was reached. One milliliter of the culture was removed as uninduced, while the remaining culture was induced using 1 mM IPTG to activate the lac promoter and induce expression of the HN and VP2 recombinant proteins. Following incubation of 4 hours, the cells were harvested by centrifugation and the pellet was re-suspended in 1:1 ratio of 1XPBS and 1X loading dye (4x dyes, 1M DTT, MilliQ water). The samples were boiled at 95°C for 10 minutes and clarified by centrifugation. The supernatant (15-20 μl) was separated on 12% SDS-PAGE with appropriate controls. Bands were observed (15-25 kDa) in the IPTG-induced lanes, which corresponds to the size of the HN and VP2 proteins respectively, and no such band was present in the Uninduced (control) lane. The recombinant proteins were batch purified using Ni – NTA column (GE Healthcare) to check the affinity of the protein with beads. Briefly, the induced culture was pelleted, and the supernatant was separated. The pellet was resuspended in 1XPBS and run on 12% SDS–PAGE. The supernatant was passed through Ni-NTA beads and the flowthrough was collected. The beads were washed with 20mM Imidazole and further eluted using 200mM Imidazole. The proteins in each sample were separated on 12% SDS PAGE and visualized by staining with Coomassie Brilliant Blue.

### Cloning, expression, and purification of rOMVs expressing antigenic fragments of HN, VP2 from NDV and IBDV, respectively

The cloned constructs for HN and VP2 in pET23a vector were used as a template for cloning both HN and VP2 in pBAD-ClyA vector separately to produce recombinant outer membrane vesicles (rOMVs). Restriction sites XbaI and HindIII were introduced at 5’and 3’ ends of HN and VP2 through forward primer and reverse primer. The polymerase chain reaction was run for 35 cycles of denaturation (30s, 95° C), annealing (30s, 59° C), and extension (25s, 72° C) in a Thermocycler (Applied Biosystems). The correctly sized amplification products were obtained for both antigens. The inserts obtained from PCR were double-digested using XbaI and HindIII restriction enzymes. pBAD-ClyA vector was also digested simultaneously using the same pair of restriction digestion enzymes to generate sticky ends compatible with the insert, and double-digested fragments were retrieved from agarose gel by using Nucleospin gel/PCR clean-up (Machery Nagel). Digested insert was ligated to vector using DNA Ligase (NEB) followed by transformation in E.coli DH5-alpha cells. Colonies were selected based on antibiotic resistance, and those colonies were inoculated for plasmid isolation. Double digestion was performed using XbaI and HindIII to confirm the clones. After digestion agarose gel electrophoresis was performed, and bands corresponding to HN and VP2 were observed for two viruses, NDV and IBDV, which confirms successful cloning of HN and VP2 in pBAD-ClyA vector.

After cloning, we further transformed confirmed clones into hypervesiculating *Ecoli* Nissle 1719 then proceeded with expression and purification of rOMVs expressing HN and VP2 according to the methods described earlier with modifications ^[8]^as well the methods we have adopted to purify rOMVs expressing GFP proteins. Vesicles were isolated by ultracentrifugation and resuspended in sterile PBS. Protein estimation of rOMVs was determined by the BCA method, and samples were loaded in an SDS gel.

### Biophysical characterization of HN/VP2 OMVs

The characterization of rOMV expressing HN/VP2 was conducted using transmission electron microscopy (TEM) (TEM-FEI Tecnai) and Dynamic Light Scattering (DLS) with equipment from Anton Paar Pvt. Ltd. For the TEM analysis, samples were suspended in water, stained with 2% uranyl acetate for two minutes, dried, and then placed on a 200-mesh type-B carbon-coated copper grid. Observations were made under a TEM, and the resulting data were collected and analyzed. DLS was employed to assess the size distribution of the OMVs in their suspension state. NTA (Nanoparticle Tracking Analysis) was also used to determine the size, charge, and number of nanoparticles in a given sample. The total protein concentration of OMVs was determined by bicinchoninic acid assay (BCA).

### Animal experiments

The University of Hyderabad’s IAEC approved the animals utilized in this research. Balb/c mice, aged 7 to 8 weeks, were orally immunized with 50 µg of soluble purified HN and VP2 proteins, as well as rOMVs expressing HN and VP2, divided into the following groups: i) PBS control; ii) soluble recombinant purified antigen (rHN and rVP2); iii) Empty OMVs (Vehicle Control); and iv) a divalent formulation of OMVs expressing HN and VP2, each mixed separately, with a 50 µg dosage of expressed antigen in the OMVs. Finally, 400 µl volume of the formulation were injected in each mouse. Blood was collected at the 15th and 28th day post-immunization to collect faeces and serum. Faeces were collected every week for IgA estimation. Mice were sacrificed after 30 days of immunization. Mice were euthanized by placing them in a chamber where 100% CO2 was introduced, displacing 30-70% of the chamber’s volume per minute and mixing with the existing air. Faeces and mucus were retrieved from the large intestine and the small intestine after being flushed with PBS, respectively.

### Localization of GFP-labelled rOMVs in the mouse intestine

Mice were orally gavaged with rOMVs expressing GFP on the surface. After some time, mice were euthanized and checked for the GFP signal by extracting the small intestine of the mouse. Further, the intestines were processed, making Swiss rolls carefully as described earlier^[28]^ using two forceps and embedded in tissue embedding media (Thermo) for sectioning using a cryotome. The sections were subsequently stained with DAPI and examined using a fluorescence microscope (Zeiss).

### Serum antibody ELISA

96 well ELISA plate (maxisorp-Nunc) was coated with 100ul/well of HN and VP2 proteins (50µg ml-1) in the bicarbonate coating buffer (pH=9.5) and incubated at 4 °C overnight as described earlier ^[29]^. Plates underwent three washes with a washing buffer (1XPBST containing 0.05% Tween-20) and were then blocked using 200µl of 4% skimmed milk for two hours at room temperature, followed by another three washes. Serum samples were diluted in 0.1% skimmed milk made with 1XPBST, added to the wells, and incubated for two hours at room temperature. After incubation, the plates were washed four times and then incubated with an HRP-conjugated anti-mouse secondary antibody for one hour at room temperature. The plates were washed five times, and 50 μl/well of TMB substrate was added. The reaction was halted at specific time points using 0.2 N H2SO4, and absorbance was measured at 450 nm.

### NDV Propagation in embryonated eggs

The NDV Komarov strain was supplied as a frozen allantoic fluid. Upon thawing on ice, the virus was diluted at a 1:10 ratio with sterile, cooled phosphate-buffered saline (PBS). The air sac was located, and a suitable injection site on the eggshell was selected and disinfected with 70% ethanol to ensure sterility of the outer surface. A small aperture was drilled approximately 0.4 cm above the edge of the air sac. Using a fine needle, 100 μl of the prepared viral solution was injected into each 9-day-old embryonated chicken egg. The puncture sites were sealed with Fevicol glue, and the eggs were placed in a specialized incubator maintained at 37°C with optimal humidity. The eggs were rotated twice daily, and after 24 hours of incubation, they were examined using an egg-candling lamp to verify embryo viability. Up to 48 hours dead embryos were immediately transferred into the refrigerator (4°C). Live embryos were returned to the incubator. The collected eggs were harvested after 12h of cooling in the refrigerator at 4°C. Then, the eggshell was cut by scissors, and the allantoic fluid was harvested using sterile syringes (unclear or bloody allantoic fluids were rejected), filtered by syringe filter (0.45μm), and stored in Eppendorf tubes. It is measured by HA immediately before pooling and filtration, where later Plaque assay is done for NDV, and it was then stored at -80°C^[30]^.

### Plaque Reduction Neutralization Assay (PRNT)

Each serum sample was diluted serially in a 2-fold manner from a starting dilution of 1:10 to 1:640 using plain DMEM (without FBS). The virus-antibody mixture was prepared by combining serially diluted serum with Newcastle Disease Virus (NDV) strain Komarov at a titer of 3 × 10^5 PFU/mL. The mixture of serum and virus was incubated at 37°C for 1 hour to facilitate neutralization. Following this, 200 µL of the mixture was added to Vero cell monolayers cultured in 48-well plates and incubated for another hour at 37°C in a 5% CO₂ atmosphere to allow the virus to adsorb. After adsorption, the cells were rinsed with 1 mL of DMEM and overlaid with 0.8% methylcellulose (Sigma, USA). The plates were then incubated for 5 days at 37°C in a 5% CO₂ environment. Post incubation, cells were fixed using 100% chilled methanol for 1 hour at 4°C, then washed with water. The fixed cells were stained with 1% crystal violet for 1 hour at room temperature and washed again to remove excess dye. Plaques were visually counted, and the neutralization titer was determined as the highest serum dilution that produced a 50% reduction in plaque count (PRNT₅₀). GraphPad Prism 8 software was used to generate neutralization curves, and nonlinear regression analysis was performed to estimate the serum dilution required to inhibit 50% and 90% of viral infection ^[31]^.

### Statistical Analysis

The data are shown as the mean ± S.E.M from three or four separate experiments. Unpaired Student’s t-test with Mann-Whitney post-test was performed using GraphPad Prism software to evaluate the significance of differences between groups. Statistical significance was set at p < 0.05.

## Results

### Expression and purification of antigenic fragments of HN and VP2 proteins

The HN protein of NDV is a key surface glycoprotein that is the major determinant of viral virulence, tropism, and host immune response ^[26]^. Similarly, VP2 is the primary structural protein of IBDV, which is highly immunogenic ^[27]^. Given the immune-dominant nature of these proteins, they were selected for the development of rOMVs based bivalent oral vaccine formulation. Since it has multiple epitopes, the highly immunogenic region of both HN and VP2 were codon optimized and synthesized to express the recombinant immunodominant antigens for the immunological studies. As these antigenic determinants are difficult to express these proteins in soluble fractions, we have used pET SUMO vector (Fig. S1A) for cloning and expression in order to express the proteins in soluble fractions. The recombinant HN and VP2 were purified successfully using Ni-NTA column chromatography (Fig. 1A and 1B).

**Fig 1.**
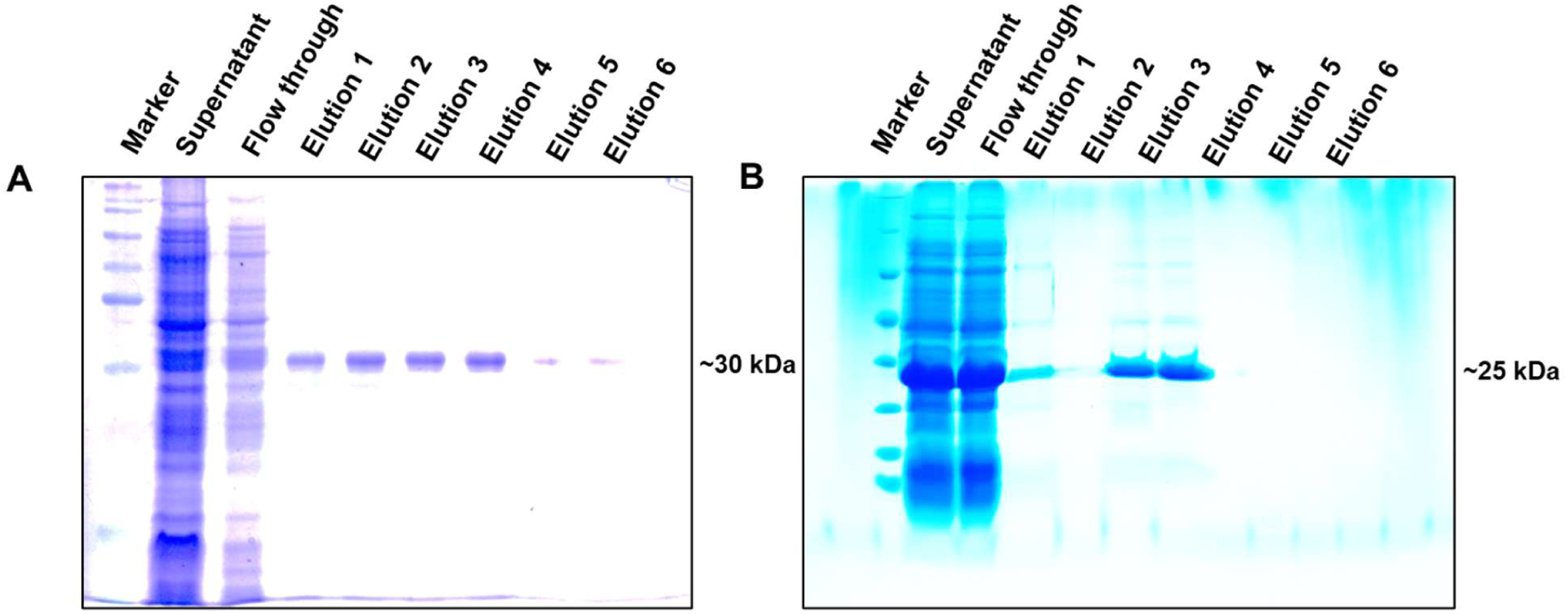
Expression and purification of soluble recombinant antigenic determinant of HN and VP2. HN cloned in pET SUMO was transformed into *E. coli* Rosetta for the expression and purification, SDS PAGE showing induction and elution fractions (E1-E6) of Recombinant HN (A). VP2 cloned in pET SUMO was transformed into *E. coli* Rosetta for the expression and purification, SDS PAGE showing induction and elution fractions (E1-E6) of Recombinant VP2 (B). Purified by Ni-NTA column chromatography.

### Expression and purification of rOMVs expressing rHN and rVP2 antigenic proteins using ClyA as a carrier partner

The strategy underlying the development of engineered nano-vesicle-based delivery systems is predicated on a fundamental principle of biology and engineering. This approach involves the fusion of two distinct proteins into a hybrid, comprising the protein of interest specifically, HN and VP2 in this study—directly linked to a membrane anchor protein, ClyA ^[8]^. ClyA is a natural constituent of the outer membrane of OMVs secreted by many bacteria, including various strains of *Escherichia coli* ^[5]^. While ClyA’s inherent function involves pore formation in membranes, we have exploited its capacity to integrate robustly with the OMV membrane. Our objective is to develop a multivalent vaccine formulation targeting poultry diseases. We have selected antigenic fragments of HN from NDV and VP2 from IBDV, successfully cloning at C-terminus of the pBAD ClyA vector. Subsequently, we confirmed, transformed, and isolated the recombinant OMVs (rOMVs) expressing HN Fig. S2 A&B) and VP2 (Fig. S2 C&D) separately. Clone confirmation was achieved through gene-specific PCR, restriction double digestion, and sanger sequencing. The expression of both ClyA HN and ClyA VP2 was verified via western blotting using an anti-His antibody (Fig 2E and F). Having successfully expressed and purified the rOMVs expressing the desired antigens, we next performed the biophysical and immunological characterization.

### Biophysical Characterization of rOMVs and localization following oral gavaging in the mice

rOMVs were purified using differential centrifugation and ultrafiltration methods as described earlier ^[10]^ with slight modifications. The size and morphology of the outer membrane vesicles were elucidated by performing Transmission Electron Microscopy (TEM) (Fig. 2A & 2B), and the distribution of the vesicles with quantification was done by Nanoparticle Tracking Analysis (NTA) (Fig. 2C & 2D) and Dynamic Light Scattering (DLS) (Fig.S3). In addition to that, we have also confirmed the presence of HN (Fig. 2E) and VP2 (Fig. 2F) expressing rOMVs by western blotting using anti-His antibody. Previously, we have demonstrated rOMVs have capability to modulate the effector functions of antigen-presenting cells (APCs), including the internalization of antigens, proteolytic processing, presentation via major histocompatibility complex (MHC) molecules, and expression of co-stimulatory molecules, which are crucial for antigen-specific adaptive immunity^[10]^. However, rOMVs and mucosal immune cell interaction were not characterized. Since the rOMVs are derived from the probiotic strain, we hypothesized that it could have mucoadhesive properties. Therefore, localization of rOMVs in the intestine was tracked by gavaging mouse with rOMVs expressing GFP. Following oral gavazing of the rOMV-GFP, the mouse was euthanized for isolating the intestine and further processed for microscopic analysis. Intestines were isolated, flushed with PBS, then made a longitudinal cut along the intestine, made a Swiss roll, and embedded in tissue freezing media for cryo-sectioning. Further slides were stained with DAPI and proceeded for fluorescence microscopy to observe the GFP signal (Fig. 2G) near the intestinal villi, where the internalization of any foreign material takes place, thereby suggesting that it could be efficiently taken by the adjoining mucosal APCs for further processing. Furthermore, its stable nature in the diverse PH conditions combined with the mucosal uptake suggests that OMVs could be tailored as an oral vaccine delivery platform

**Fig 2.**
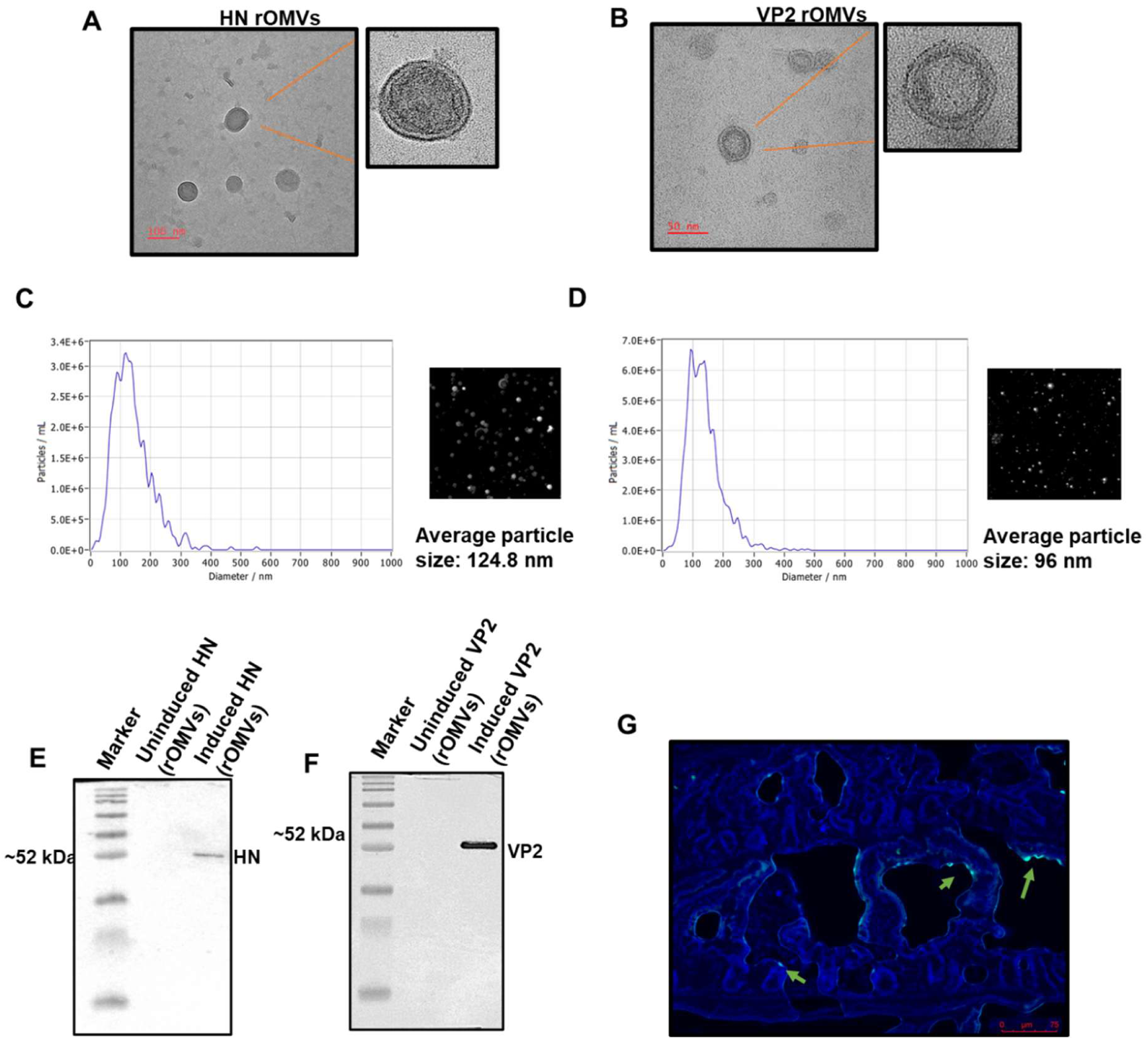
Biophysical characterization and localization of rOMVs expressing HN and VP2 antigens. rOMVs secreted by pBAD-ClyA-HN-His and pBAD-ClyA-VP2-His vectors transformed *Ecoli* Nissle 1917 cells were isolated separately and characterized for their size and approximate particle counts using Transmission Electron Microscopy (TEM) (A and B) and the Nano Particle Tracking Analysis method (NTA) (C and D) below. The further presence of HN and VP2 expressed on OMVs was confirmed by Western blotting using an anti-His antibody (E and F), respectively. To confirm whether OMVs can cross the intestine, we have cloned GFP into pBAD-ClyA and expressed it in OMVs for tracking—intestinal cross-section of a mouse gavaged with rOMV-GFP showing GFP signal (G).

### rOMVs expressing HN and VP2 elicit a robust mucosal and systemic immune response upon oral immunization

After successful expression and characterization of rOMVs, we have proceeded with the oral immunization because it is inexpensive, needle-free, and reduces costs and manpower thereby opening new avenue for the development of new generation biotherapeutics formulations, especially vaccines. Outer membrane vesicles (OMVs) can be engineered to express desired proteins, thereby serving not only as a delivery mechanism but also as an immunomodulator due to the presence of pathogen-associated molecular patterns (PAMPs) thereby sufficing the combinatorial problem of adjuvanting and oral antigen delivery. ^[9]^ ^[22]^. Therefore, to examine the potential of rOMVs in tailoring mucosal protective immunity, the mice were, orally gavaged with rOMVs expressing HN and VP2 as a cocktail (Divalent) and further evaluated the immune responses in comparison to soluble antigens alone and controls as per the immunization schedules as described in Fig 3A. Mice were sacrificed after 30 days, and indirect ELISA analyzed mucosal antigen-specific immune responses. The results showed that the antigen expressed in rOMVs induced significantly higher antibody responses, particularly sIgA levels in both fecal and mucus samples (Fig. 3 B-E), when compared to the control groups. We have also assessed the IgG levels in the serum isolated from mice immunized with rOMV vaccine formulations, where we observed strong systemic responses (Fig.4 A & B) through the mucosal-mediated immune responses.

**Fig 3.**
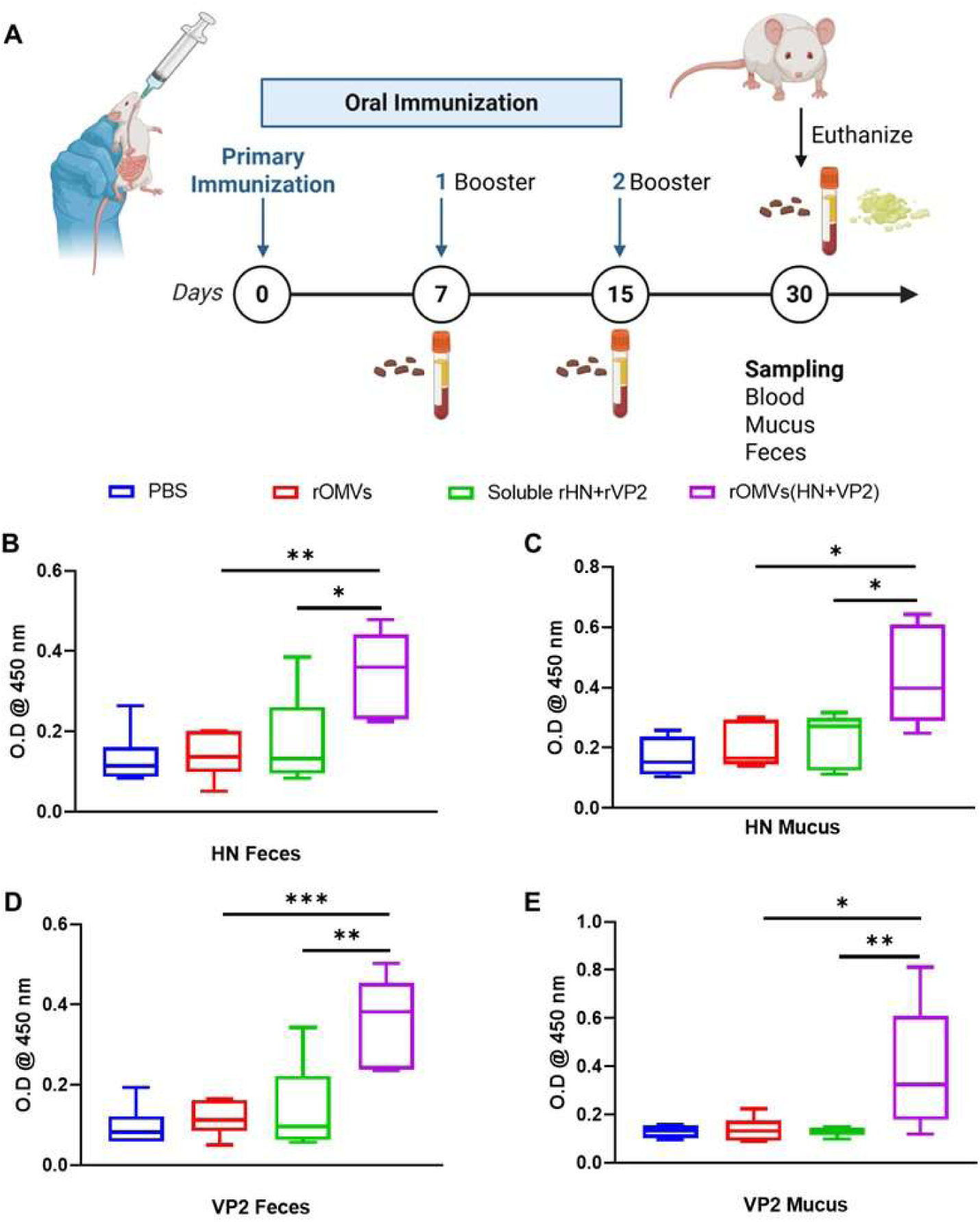
rOMV expressing HN and VP2 from NDV and IBDV viruses in a divalent formulation induces a significant antigen-specific antibody response (sIgA). A) Immunization Schedule B, C, D&E) Mice immunized orally with rOMVs expressing HN and VP2, ELISA showing sIgA from both Feces and Mucus isolated after immunization. Data of each group contains 6-7 animals, and data are represented as mean with SEM. Value of significance *P < 0.05, **P < 0.005, and ***P < 0.0005.

**Fig 4.**
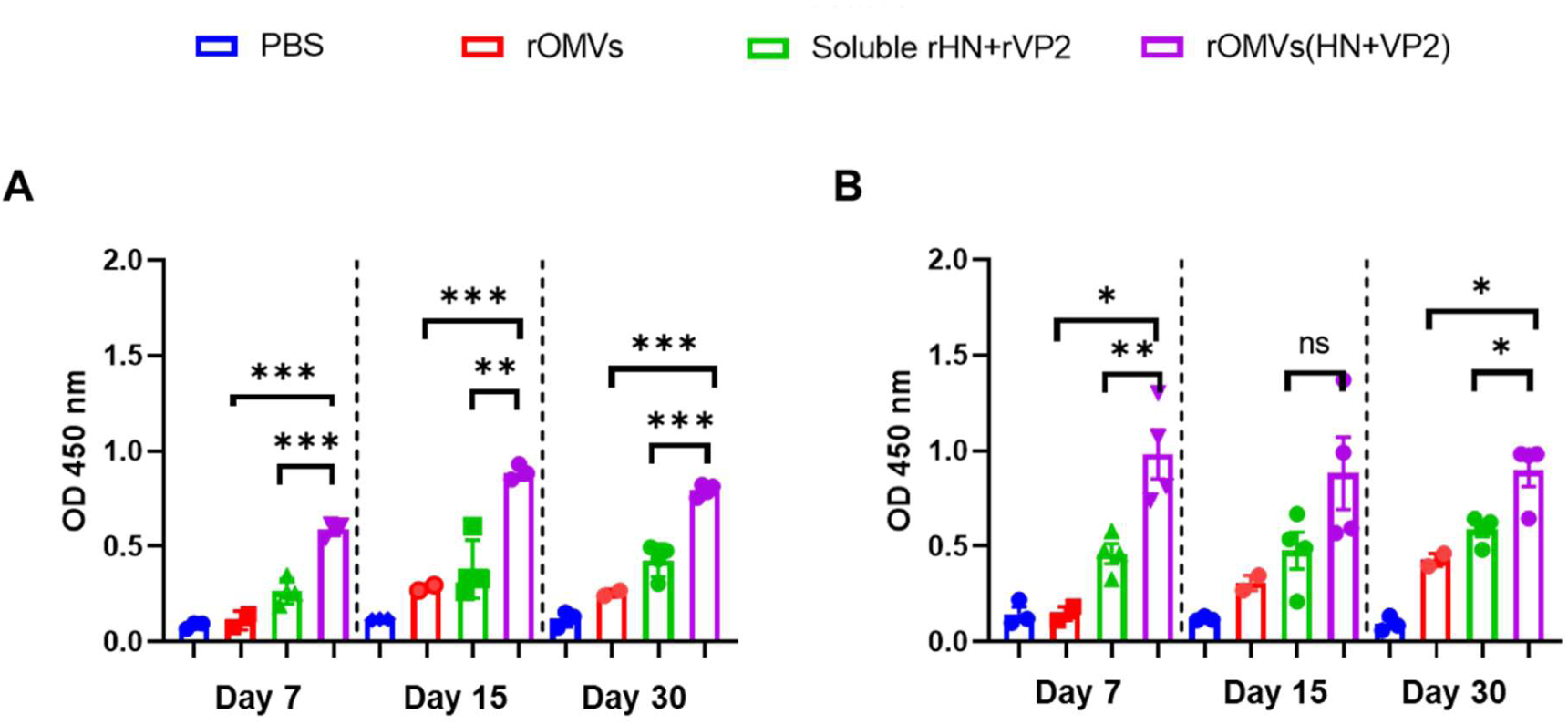
Oral administration of rOMV containing HN and VP2 from NDV and IBDV viruses in a dual formulation triggers a notable systemic antibody response specific to the antigens. IgG response in serum from mice after oral immunization, rHN-specific IgG (A) and rVP2-specific IgG (B). Data of each group contains 4-5 animals, and the data are shown as the mean along with the SEM. Significance levels are indicated as P < 0.05, *P < 0.005, and ***P < 0.0005.

### Antibodies generated after oral immunization were capable for neutralizing the virus in an invitro infection system

The potency of any vaccine delivery or adjuvanting systems lies in its ability to program the protective vaccine immunity, including high-titer neutralizing antibodies capable of neutralizing the pathogen. Here, we utilized invitro plaque reduction neutralization test to evaluate the potency of the antibodies that are generated upon oral immunizations of the rOMVs-based vaccine formulation. We have assessed the virus-neutralization capability of serum antibodies from orally immunized mice using an *in vitro* infection model. Newcastle Disease virus (NDV Komarov strain) was supplied as a frozen allantoic fluid. Virus was further grown in chicken eggs and then allantoic fluid was collected, filtered, confirmed with HA immediately and stored at -80 for plaque assays. Serum samples were serially diluted in a 2-fold dilution factor from a starting dilution of 1:10 to 1:640 using incomplete DMEM (without FBS). The virus-antibody mixture was prepared by combining serially diluted serum with Newcastle Disease Virus (NDV) strain Komarov at a titer of 3 × 10^5 PFU/mL. After incubation of serum with virus, we have proceeded for staining and counting. With increasing serum dilution number of plaques were also increasing. The neutralization titer was defined as the highest dilution of serum that produced a 50% reduction in the number of viral plaques, referred to as the PRNT₅₀ value. Neutralization curves were generated using GraphPad Prism 8 software, and nonlinear regression analysis was performed to estimate the serum dilution required to inhibit 50% and 90% of viral infection. Based on the results obtained from PRNT as compared to the control groups, rOMVs expressing HN group showed significant neutralization capacity (Fig.5), thereby suggesting the rOMV based oral vaccine formulation has the ability to tailor the mucosal immunity by triggering high titer antibodies which are capable of neutralizing the virus effectively. These data validate the rOMV systems holds considerable potential for oral vaccine formulations

**Fig 5.**
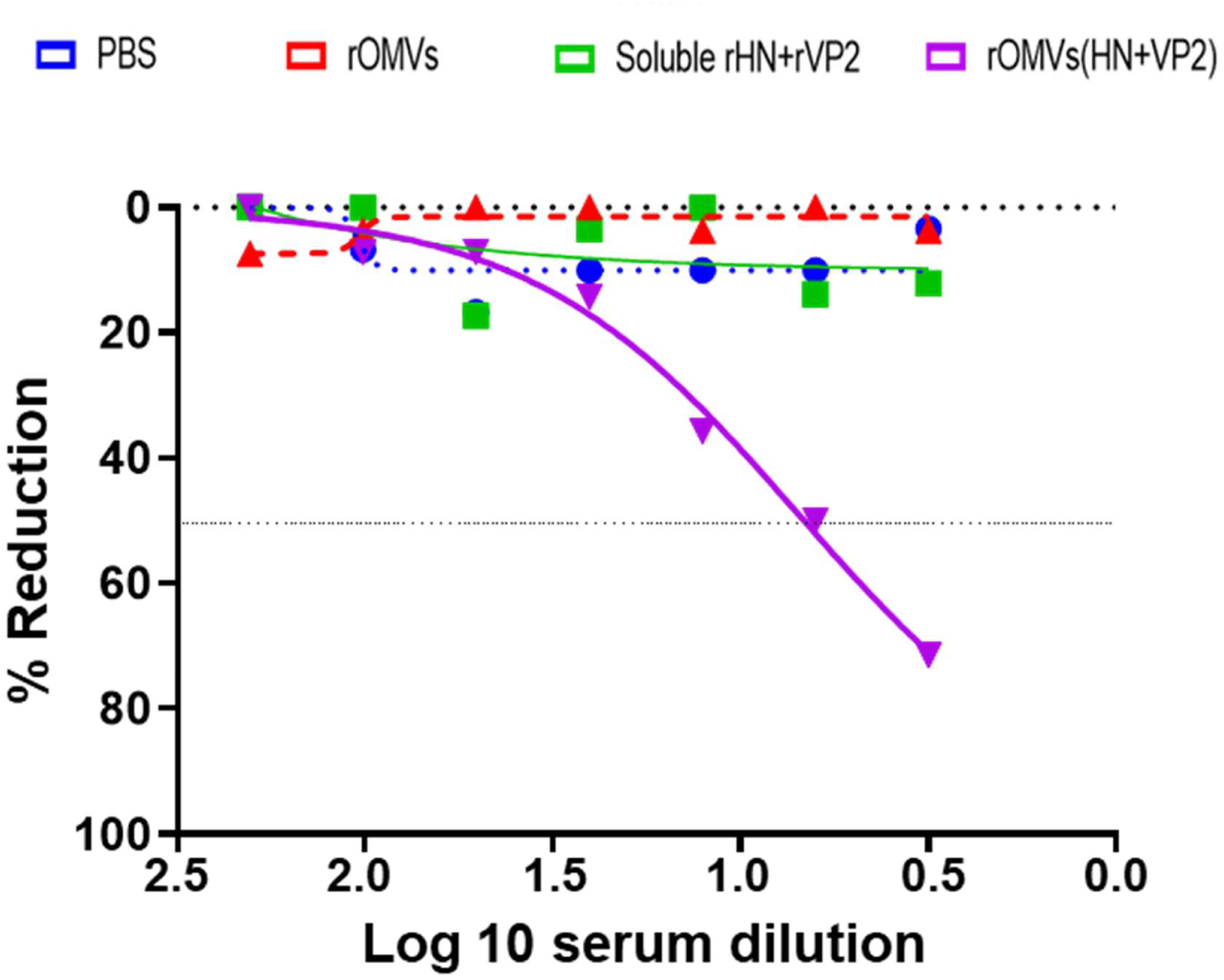
Serum from mice immunized with rOMVs expressing HN antigen effectively neutralizes Newcastle Disease Virus (NDV). The ability of immunized serum to neutralize the Newcastle Disease virus in vitro was evaluated at different dilutions using the Plaque Reduction Neutralization Test (PRNT). The PRNT50 value was determined for each immunized serum sample, which indicates the serum dilution required to neutralize 50% of the virus, as determined by the PRNT neutralization assay for NDV. The data is presented as the mean of multiple biological and experimental replicates, along with the standard deviation, and a significance level of p < 0.05 was considered significant, analyzed by student t-test followed by the Mann-Whitney test.

## Discussion

While vaccination plays a crucial role in managing poultry health, there is a need for effective, low-cost, scalable, and easily administered new-generation vaccine discovery platforms to tackle emerging infectious diseases ^[18]^. Oral vaccines present an alternative to injectable ones due to their simplicity of administration; however, current platforms do not provide strong protective systemic and mucosal immunity ^[32]^ ^[35]^. Therefore, development of cost-effective, vaccines is a urgent requirement for controlling Poultry infectious diseases such as Newcastle Disease Virus (NDV) and Infectious Bursal Disease Virus (IBDV), whose outbreaks continue to pose major economic and animal welfare concerns ^[33]^. In the current study, we successfully demonstrated the expression, purification, surface display, and immunogenicity of recombinant antigen-loaded outer membrane vesicles (rOMVs) as a potential oral mucosal vaccine platform targeting these two important avian pathogens. By utilizing a novel bioengineering strategy that combines recombinant protein technology with outer membrane vesicle biology, we provide compelling evidence that this platform could serve as a promising oral delivery alternative to traditional vaccine approaches.

The expression of viral proteins such as the hemagglutinin-neuraminidase (HN) from NDV and the viral protein VP2 from IBDV in heterologous bacterial systems has historically been challenging, primarily due to issues related to protein solubility and folding. To address this, we employed codon optimization techniques to adapt the antigen sequences for efficient expression in *Escherichia coli*, followed by cloning into the pET-SUMO vector. The choice of the SUMO (Small Ubiquitin-like Modifier) tag was strategic, as it is well-documented to enhance the solubility and stability of difficult-to-express proteins in bacterial systems ^[34]^. Using Ni-NTA affinity chromatography, both recombinant proteins were successfully purified, establishing a foundation for downstream antigen engineering.

Subsequent efforts focused on the generation of rOMVs expressing these viral antigens. To achieve this, we adopted a membrane-anchoring strategy using the cytolysin A (ClyA) protein, a naturally occurring component of the bacterial outer membrane ^[34]^. ClyA is known for its strong integration into the outer membrane and its ability to tolerate large C-terminal fusions without compromising membrane localization. By genetically fusing the HN and VP2 antigens to the C-terminus of ClyA and expressing these constructs under the arabinose-inducible pBAD promoter in *E. coli* Nissle, we effectively generated OMVs decorated with the desired antigens.

A critical advantage of mucosal vaccines lies in their ability to target immune-inductive sites directly at the portal of pathogen entry ^[11]^. To assess whether the rOMVs could localize to the intestinal mucosa following oral administration, we used a green fluorescent protein (GFP)-tagged OMV construct. Mice were gavaged with GFP-rOMVs, and intestinal tissues were collected and processed for fluorescence microscopy. GFP fluorescence was observed near the villi of the small intestine, indicating the successful localization and potential interaction of the rOMVs with mucosal immune components. This observation is particularly important, as the uptake of antigens at mucosal surfaces initiates immune responses that can prevent pathogen colonization and dissemination at their primary entry points. Moreover, this would aid in tailoring the mucosal APC in programming the protective immunity.

Having observed that the rOMVs are effectively localized in the mucosal sites, next; we evaluated the immunogenicity of the rOMVs expressing HN and VP2 upon oral administration in mice. Oral immunization offers multiple logistical advantages elimination of needles, reduction in labor costs, and better compliance and is particularly attractive for veterinary settings where mass immunization is needed. In our study, mice were orally immunized with a cocktail formulation containing both rOMVs-HN and rOMVs-VP2. Analysis of fecal and intestinal mucus samples revealed significantly elevated levels of antigen-specific IgA, an essential mucosal antibody that plays a pivotal role in pathogen neutralization at mucosal surfaces. Compared to mice that received soluble antigen alone or control treatments, the rOMV-immunized groups exhibited markedly higher mucosal IgA levels, indicating a successful priming of the mucosal immune system.

In addition to mucosal responses, serum IgG levels were also analyzed to determine the systemic immune activation mediated by the rOMV vaccine platform. While the IgG titers were comparatively lower than the IgA levels, they were nonetheless detectable, suggesting a degree of systemic dissemination of the antigen or antigen-presenting cells. The dual induction of both mucosal and systemic responses is desirable in vaccine development, particularly for pathogens like NDV and IBDV that can breach mucosal barriers and induce systemic infections.

To functionally validate the protective potential of the antibodies elicited by the rOMV platform, we conducted a plaque reduction neutralization test (PRNT) using the NDV Komarov strain. PRNT is considered the gold standard for assessing virus-neutralizing antibodies, as it directly measures the ability of serum antibodies to prevent viral infection in a permissive cell culture model. Our results demonstrated that sera from rOMV-HN immunized mice showed a significant reduction in viral plaque formation in a dose-dependent manner. The neutralizing activity was quantified as PRNT50 values, which reflect the serum dilution at which 50% of the viral plaques are inhibited. The observed PRNT50 titers from the rOMV-HN group indicate that the generated antibodies are not only specific but functionally capable of neutralizing the virus, a critical requirement for protective immunity. This is a strong indicator of the potential of the rOMV system to serve as a viable NDV vaccine candidate.

The results presented here align well with the growing body of literature highlighting OMVs as a potent vaccine delivery platform. Their intrinsic immunostimulatory properties, derived from components such as lipopolysaccharide (LPS), outer membrane proteins, and peptidoglycan, render OMVs inherently adjuvant-like. Importantly, the LPS levels in OMVs derived from *E. coli* Nissle 1917 are lower and less reactogenic compared to pathogenic strains, making them suitable for use in oral formulations. Furthermore, OMVs possess a natural tropism for uptake by antigen-presenting cells, facilitating efficient antigen processing and presentation.

Taken together, this study provides a compelling proof-of-concept for the use of bioengineered OMVs as an oral vaccine delivery system targeting NDV and IBDV. The combination of high antigen-specific mucosal responses, demonstrable virus neutralization activity, and scalable production in a probiotic *E. coli* chassis makes this platform especially promising for poultry vaccination programs. Given the limitations of current live-attenuated or inactivated vaccines, such as cold chain dependency, injection-based administration, and potential reversion to virulence, the use of non-replicative, recombinant OMVs offers a safer and more versatile alternative. Future directions for this work include the evaluation of protective efficacy in avian challenge models, longitudinal analysis of immune memory, and optimization of the antigen payload and dosing regimens. Additionally, co-display of multiple antigens on a single OMV, or the use of tandem OMV cocktails, could be explored to generate broad-spectrum protection against multiple serotypes or co-circulating avian pathogens. Further safety assessments and regulatory considerations will also be crucial before field deployment.

In conclusion, this study demonstrates that engineered OMVs expressing viral antigens HN and VP2 can serve as an effective and immunogenic oral vaccine platform. The data presented support the potential of this technology to address current gaps in poultry vaccine delivery, offering a needle-free, scalable, and immunologically robust alternative to conventional vaccines. As the field of OMV-based vaccinology continues to evolve, our findings contribute meaningful insights into the practical and translational prospects of mucosal immunization strategies in veterinary and potentially human applications.

## Conclusion

The widespread occurrence of infectious diseases, such as IBD, IBV, and ND, has significantly hindered the growth of the poultry industry for many years. Existing vaccines for these diseases have limitations such as poor effectiveness, limited immune response, and adverse reactions. To address these issues, this study explored the use of outer membrane vesicles (OMVs) as an oral vaccine delivery vehicle as well as a natural adjuvant system. OMVs can improve the immunogenicity and sustained release of key viral proteins, such as HN and VP2 proteins of NDV and IBDV. These proteins have been identified as potential vaccine candidates for enhancing the immunity against infectious diseases. These key findings from our study showed the use of rOMVs in delivering vaccine antigens effectively, safely, and economically feasible for tackling poultry diseases, which could be extrapolated to human diseases.

## Author Contributions

**SD:** Investigation, designed and performed experiments, analyzed data, and Manuscript writing (original draft). **FA:** Manuscript editing and experimental design. **RAK:** Assisted in the initial design of experiments.**SN**: helped with animal experiments, **SNC:** Bioinformatics, **MS:** Recombinant protein purification. **AM:** Helped with protein purification. **TJ**: Performed plaque assay. **MS:** Helped with NDV viral assay and review**. NK** developed the research concept, oversaw the project, and assisted with manuscript revisions and securing funding. All authors have reviewed and approved the final version.

## Acknowledgments

The Authors would like to thank all the facilities provided by the School of Life Sciences, University of Hyderabad, including the flow cytometry and Transmission Electron Microscopy core facilities

## Ethical statement

The experiments were conducted in line with protocols sanctioned by the University of Hyderabad’s institutional biosafety committees.

## Data availability

This research did not utilize data from external sources. All pertinent data are contained within the manuscript and the accompanying supplementary files.

---

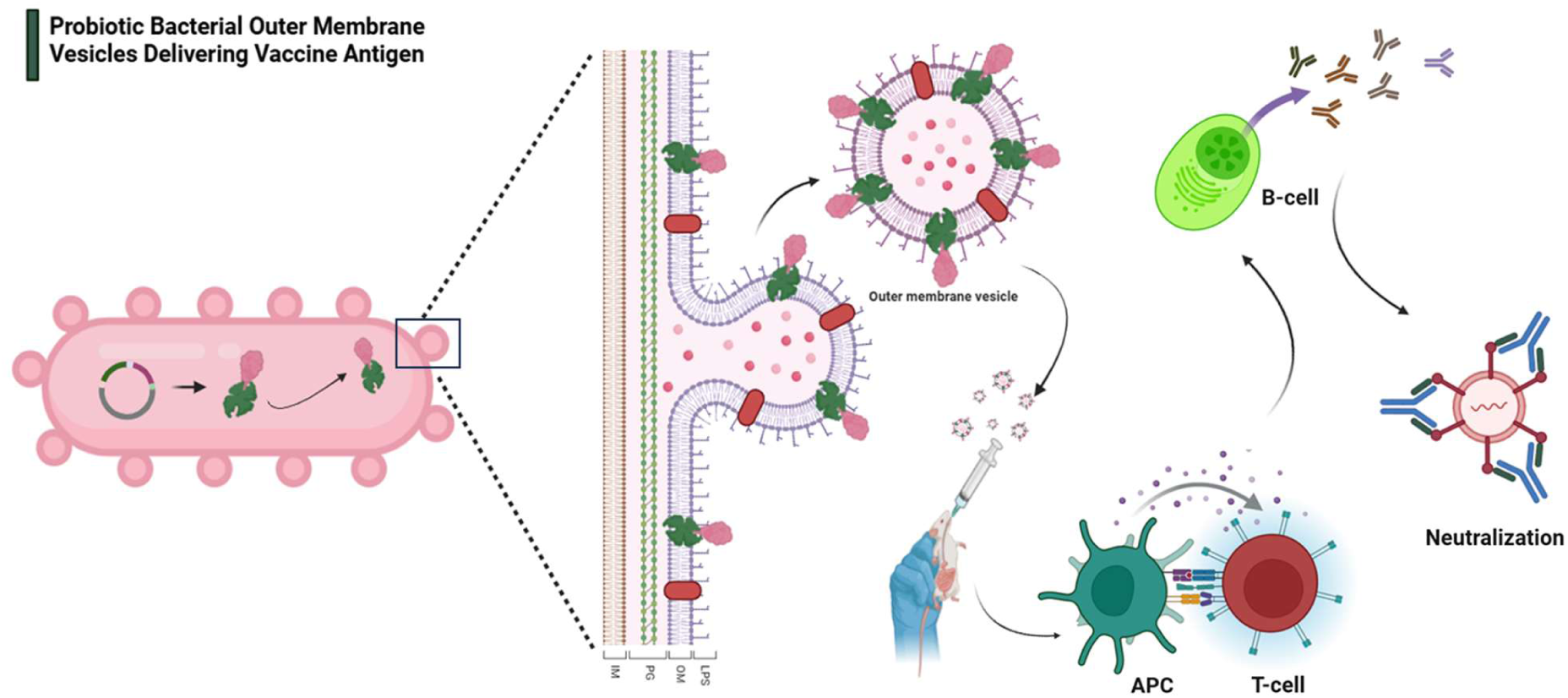

