## Supplementary Material for "Oral administration of engineered recombinant outer membrane vesicles-based nano-vaccine formulation triggers robust mucosal and systemic humoral immunity against poultry pathogens"

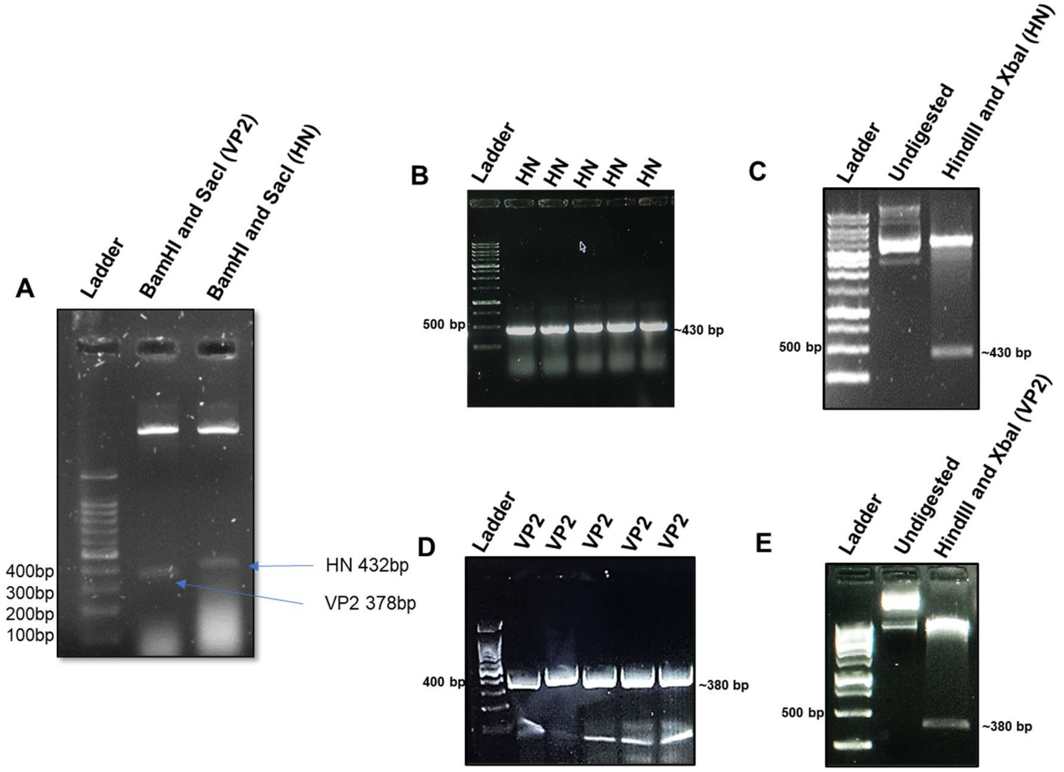


**Figure S1.** Clone confirmation of HN/VP2 in pBAD-ClyA vector.. Synthesized clones were confirmed by double digestion (A). The HN gene was cloned into pBAD-ClyA-His Vector and transformed into Nissle Ecoli 1917 cells, clone confirmation by colony PCR (gene specific primers) (B) with HN-specific primers, double digestion by Xbal and Hindlll (C). The VP2 gene was cloned into pBAD-ClyA-His Vector and transformed into Nissle Ecoli 1917 cells, clone confirmation by colony PCR (D) with HN-specific primers, double digestion by Xbal and Hindlll (E).

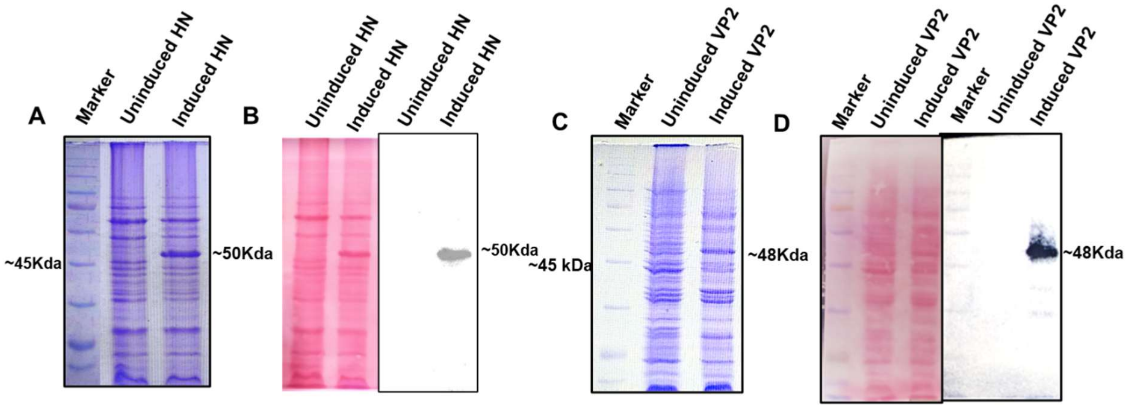


**Fig S2.** Expression and purification of HN/VP2 in *Ecoli* Nissle 1917. Expression of ClyA-HN in *Ecoli* Nissle 1917 induced with 0.2 % arabinose and confirmed using Coomassie (A), Immunoblotting and representative ponceau S staining (B). Purified by Ni-NTA column chromatography. Expression of ClyA-VP2 in Nissle Ecoli induced with 0.2 %arabinose and confirmed using Coomassie (C), Immunoblotting and representative ponceau S staining(D).

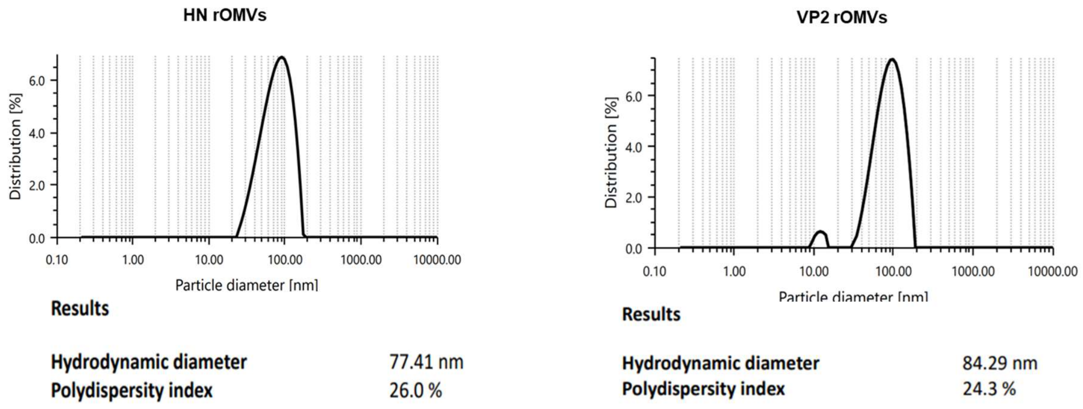


**Figure S3**: Biophysical characterization A&B) Dynamic Light Scattering (DLS) to show the size distribution of HN-OMVs and VP2-OMVs.
